# Distinguishing Happiness and Meaning in Life from Depressive Symptoms: a GWAS-by-subtraction study in the UK Biobank

**DOI:** 10.1101/2022.12.06.519260

**Authors:** Lianne P. de Vries, Perline A. Demange, Bart M.L. Baselmans, Christiaan H. Vinkers, Dirk H.M. Pelt, Meike Bartels

## Abstract

**Background:** Hedonic (e.g., happiness) and eudaimonic (e.g., meaning in life) well-being are negatively related to depressive symptoms. Genetic variants play a role in this association, reflected in substantial genetic correlations. We investigated the (genetic) overlap and differences between well-being and depressive symptoms.

**Methods:** We used results of Genome-Wide Association studies (GWAS) and applied GWAS-by-subtraction in the UK Biobank sample. Analyses were pre-registered.

**Results:** Subtracting GWAS summary statistics of depressive symptoms from those of happiness and meaning in life, we obtained GWASs of respectively ‘pure’ happiness (n_effective_= 216,497) and ‘pure’ meaning” (n_effective_=102,300). For both, we identified one genome-wide significant SNP (rs1078141 and rs79520962, respectively). After the subtraction, SNP heritability reduced from 6.3% to 3.3% for pure happiness and from 6.2% to 4.2% for pure meaning. The genetic correlation between the well-being measures reduced from .78 to .65, indicating that only a part of the genetic overlap between happiness and meaning in life is due to overlap with depressive symptoms. Pure happiness and pure meaning became genetically unrelated to traits strongly associated with depressive symptoms, including tiredness, loneliness, and psychiatric disorders. For several other traits, including ADHD, income, educational attainment, smoking, and drinking alcohol, the genetic correlations of well-being versus pure well-being changed substantially.

**Conclusions:** GWAS-by-subtraction allowed us to investigate the genetic variance of well-being unrelated to depressive symptoms. Genetic correlations with different traits led to new insights about this unique part of well-being. The findings can have implications for interventions to increase well-being and/or decrease depressive symptoms.

## Introduction

In the past, well-being and ill-being, such as depressive symptoms, have been considered opposite ends of a continuum. However, the overlap between well-being and depressive symptoms is only moderate. Phenotypic correlations range between −.40 and −.60 (1–3) and genetic correlations from −.50 to −.81 (1,4,5). Well-being and ill-being are thus seen as distinct, but related domains of mental health.

A distinction is often made between hedonic well-being and eudaimonic well-being (6). Hedonistic philosophical ideas on well-being include maximizing pleasure and minimizing pain (6,7). Modern-day hedonic well-being measures focus on levels of positive and negative affect, and satisfaction with life (8). Eudaimonic philosophical theories extend beyond pleasure and pain, and emphasizes living a virtuous life (6,7). Current eudaimonic well-being measures include measures of positive functioning, thriving, and judgments about the meaning and purpose of life (9). In this project we operationalized hedonic well-being with a measure of happiness and eudaimonic well-being with a measure of meaning in life.

Hedonic and eudaimonic measures of well-being have been found to load on separate, but correlated factors (>.60) (10–12). In recent molecular genetic work, the moderate phenotypic correlation between happiness (hedonic well-being) and meaning in life (eudaimonic wellbeing) was replicated (r_ph_ = 0.53) and a strong genetic correlation (r_g_ = 0.78) was observed, suggesting a largely shared genetic etiology (5). Furthermore, genetic correlations with related traits were similar for happiness and meaning in life (5). The only genetic correlation that differed for happiness compared to meaning in life was with depressive symptoms, with a moderate genetic correlation for happiness (r_g_ = −0.53, *SE*=.04), and a smaller correlation for meaning in life (r_g_ = −0.32, *SE*=.05).

The reported phenotypic and genetic correlations between happiness, meaning in life and depressive symptoms indicate substantial overlap between well-being and depressive symptoms. However, less is known about the part that makes well-being unique, i.e., independent from depressive symptoms. Recently, GWAS-by-subtraction was developed in order to disentangle the shared and unique genetic variance for traits (13). To further investigate the (genetic) overlap and differences between happiness and meaning in life, and the overlap with depressive symptoms, we applied GWAS-by-subtraction on UK Biobank data. Subtracting a depressive symptoms GWAS from happiness and meaning GWASs in UK Biobank (5), we obtained GWASs of respectively ‘pure’ happiness and ‘pure’ meaning. In follow-up analyses, we investigated the genetic variants associated with pure happiness and pure meaning using functional annotation and genetic correlations with other traits.

## Methods and Material

### Participants

UK Biobank is a large, population-based prospective study with data from over half a million participants of middle to old age from the United Kingdom (14). During the initial assessment visit (2006-2010) a touchscreen questionnaire was used to collect extensive information, including sociodemographic characteristics, lifestyle exposures and general health from the participants. In a later follow-up (2016), participants completed online questionnaires, including mental health and well-being questions.

We used data from 427,580 participants with genetic data and data on depressive symptoms from the initial assessment. Permission to access both phenotypic and genetic UK Biobank data was obtained under application number 40310. Furthermore, we used the summary statistics of Baselmans and Bartels (2018) on happiness (n= ~222k individuals) and meaning in life (n= 108k individuals) in UK Biobank participants.

### Depressive symptoms

In line with Okbay et al. (2016), to create a depressive symptoms score, we summed standardized scores on two items; *Over the past two weeks, how often have you felt down, depressed or hopeless?* (UKB Data-Field 2050), and *Over the past two weeks, how often have you had little interest or pleasure in doing things?* (UKB Data-Field 2060). Participants answered on a 4-item Likert scale that ranged from “Not at all” (1) to “Nearly every day” (4).

### Genetic data

Genome-wide genotype data for the participants have been collected, processed, quality controlled and imputed by UK Biobank (see for a full description Bycroft et al., 2018). To briefly summarize, participants were assayed using two similar genotyping arrays, the Affymetrix UK BiLEVE and UK Biobank Axiom Arrays. The phasing and imputing were performed using the Haplotype Reference Consortium and merged UK10K and 1000 Genomes phase 3 reference panels. The quality control was designed to address issues specific to a large-scale dataset. Quality control steps for markers included testing for batch effects, plate effects, departures from Hardy–Weinberg equilibrium, sex effects, array effects, and discordance across control replicates. Samples were excluded based on non-European ancestry, sex mismatch between genetic result and self-report, and metrics of missing rate and heterozygosity (15).

### Statistical analyses

The analyses were pre-registered before data analysis (https://osf.io/pnc2z).

#### GWAS depressive symptoms

The genome-wide association analysis on the created depressive symptoms score was performed in GCTA using linear mixed modelling (LMM). This controls for population stratification by including a genetic relatedness matrix (GRM) (16). As additional controls, we included sex, age, sex*age, and 100 genetic principal components. We used the recommended threshold of *p* < 5×10^-8^ for significant SNPs and *p* < 1×10^-5^ for possible implicated SNPs (17).

#### GWAS-by-subtraction

We used Genomic Structural Equation Modelling (Genomic SEM) (18) and GWAS-by-subtraction (13) to investigate the overlap between depressive symptoms and respectively happiness and meaning in life. For each SNP, GWAS-by-subtraction estimates the association with a trait of interest that is independent of the association of that SNP with another trait, in our case well-being and depressive symptoms. In the model, the GWAS summary statistics of both traits are regressed on two latent variables, i.e., *Depressive Symptoms* and *Pure Happiness* or *Pure Meaning* (see lower part of Figure 1). These latent factors are regressed on each SNP (see top part of Figure 1). For each SNP, this model results in two paths of association. In one path, the SNP effects are mediated by depressive symptoms. The other path is independent from depressive symptoms and indicates the SNP effects for pure well-being. In other words, the variance of well-being is separated in a part shared between well-being and depressive symptoms, and in a part unique for well-being, i.e., pure well-being. The genetic variance for pure happiness and pure meaning is by design independent of the genetic variance for depressive symptoms (r_g_ = 0). We performed sensitivity analyses to investigate the impact of a bidirectional effect (see supplementary Material).

**Figure 1.**
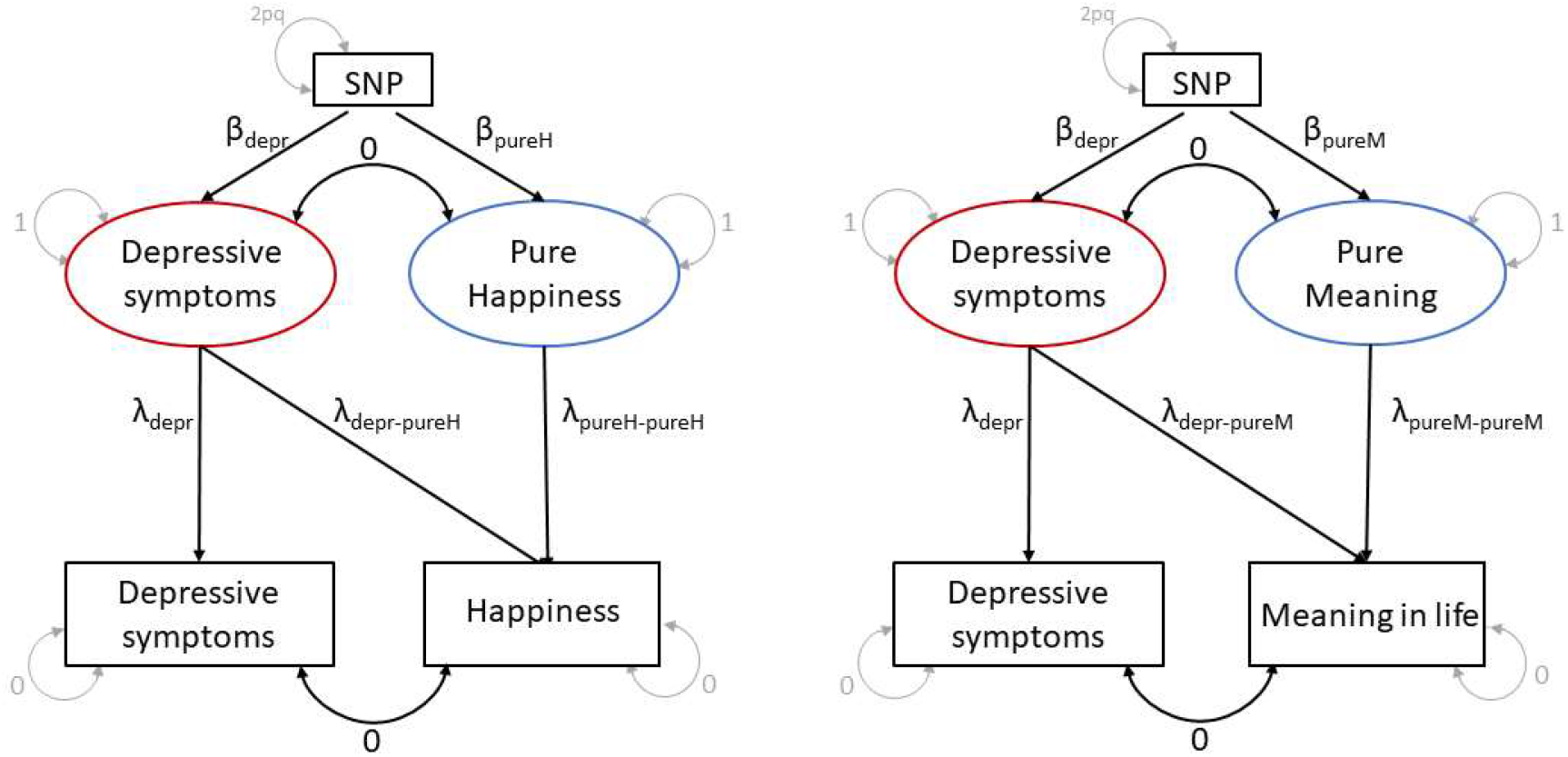
Schematic overview of the GWAS-by-subtraction approach to create a GWAS of “pure happiness” and “pure meaning”.

#### GWAS follow-up analyses

##### SNP heritability

Univariate and bivariate LD score regression (LDSC) (19) was used to estimate the SNP heritability for pure happiness and pure meaning and to compute the genetic correlation between happiness and pure happiness and between meaning in life and pure meaning.

##### Functional annotation

For pure happiness and pure meaning, we looked up the lead significant SNPs (*p* < 5×10^-8^) in the NHGRI-EBI catalogue of human genome-wide association studies (www.ebi.ac.uk/gwas/).

To follow-up on the SNP based association test for pure happiness and pure meaning in life, we performed gene mapping in FUMA (http://fumactglab.nl, (20)). Gene mapping was based on three strategies, namely positional mapping (i.e., physical distance from the gene, within 10 kb window), eQTL mapping (i.e., the gene expression is associated with allelic variation at the SNP), and chromatin interaction mapping. Furthermore, we applied genome-wide gene-based association tests using MAGMA (21). The gene-based test combines results from multiple SNPs within a gene to test the association between genes and pure happiness or pure meaning, while accounting for LD between SNPs.

#### Genetic correlations

To further investigate the distinction between pure happiness, pure meaning, and depressive symptoms, we calculated genetic correlations between these traits and a range of other traits, using bivariate LDSC regression. We included selected traits across 12 categories with well-powered GWAS data (N=75 GWAS, see supplementary Table 4) and used a Bonferroni corrected threshold (*p* = 0.05/(75*5) = 1.3×10^-5^).

## Results

### GWAS depressive symptoms

A depressive symptoms score was computed for 467,389 participants (M= 2.58, *SD* = 1.12, range = 2-8). 427,580 individuals had genetic data available and were included in the GWAS. The depression GWAS resulted in 14 independent genome-wide significant SNPs (λ_GC_ = 1.32, LD intercept = 1.02) and a SNP heritability of 4.4% (*SE*=0.002). The results and Manhattan plot can be found in supplementary Table S1 and Figure S1.

### GWAS-by-subtraction depressive symptoms and happiness

GWAS-by-subtraction of depressive symptoms and happiness resulted in one independent genome-wide significant SNP for pure happiness (N_effective_= 216,497) (λ_GC_ = 1.13, LD intercept = 0.99). The significant SNP was rs1078141 (CHR:BP = 8:142619393, *β* = 0.102, *SE*=0.018, *Z* = 5.73, *p* = 1.03×10^-8^). The results from the pure happiness GWAS are shown in the Manhattan plot in Figure 2 and the QQ plot in supplementary Figure S2. SNPs for happiness (5) and pure happiness are compared in supplementary Table S2.

**Figure 2.**
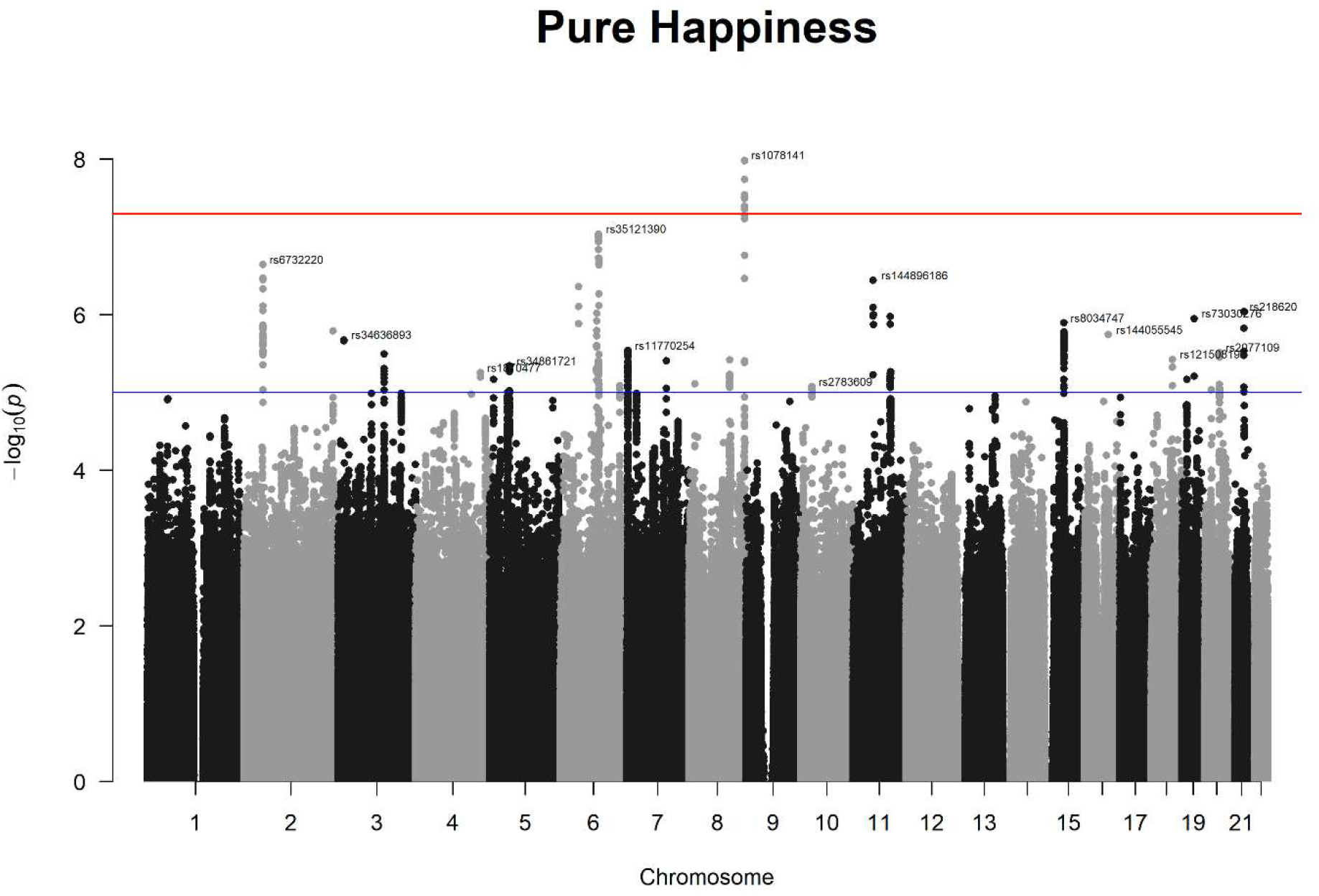
Manhattan plot for the GWAS results of pure happiness.

#### SNP heritability and genetic correlation

The SNP heritability of pure happiness was estimated to be 3.3% (*SE* = 0.003), a reduction of ~3% compared to the SNP h^2^ of 6.3% (*SE* = 0.005) for happiness in Baselmans and Bartels (2018). The genetic correlation between pure happiness and happiness was 0.80 (*SE* = .02, *Z* = 52.51, *p* < .001), indicating a reduction in genetic (co)variance.

#### Functional annotation

The effect of the significant SNP rs1078141 of pure happiness (*β* = .102, *p* = 1.03×10^-8^) is similar to the effect of this SNP in Baselmans and Bartels (2018) (*β* = .017, *p=* 5.57×10^-8^). The look-up showed that the significant SNP has also been associated with general cognitive ability (22).

Applying FUMA, no genes were associated with pure happiness based on positional mapping, eQTL mapping, or chromatine interaction-mapping. The gene-based test indicated no significant genes, and no significant enrichment of genes in certain tissues was found for pure happiness.

### GWAS-by-subtraction Depressive Symptoms and Meaning

The GWAS-by-subtraction of depressive symptoms and meaning resulted in 1 genomewide significant SNP for pure meaning (N_effective_=102,300) (λ_GC_ = 1.08, LD intercept = 0.99). The significant SNP was rs79520962 (CHR:BP = 7:127671511, *β* = 0.304, *SE* = 0.054, *Z* = 5.62, *p* = 1.86×10^-8^). The results from the pure meaning GWAS are shown in the Manhattan plot in Figure 3 and the QQ plot in supplementary Figure S3. SNPs for meaning in life (5) and pure meaning are compared in supplementary Table S2.

**Figure 3.**
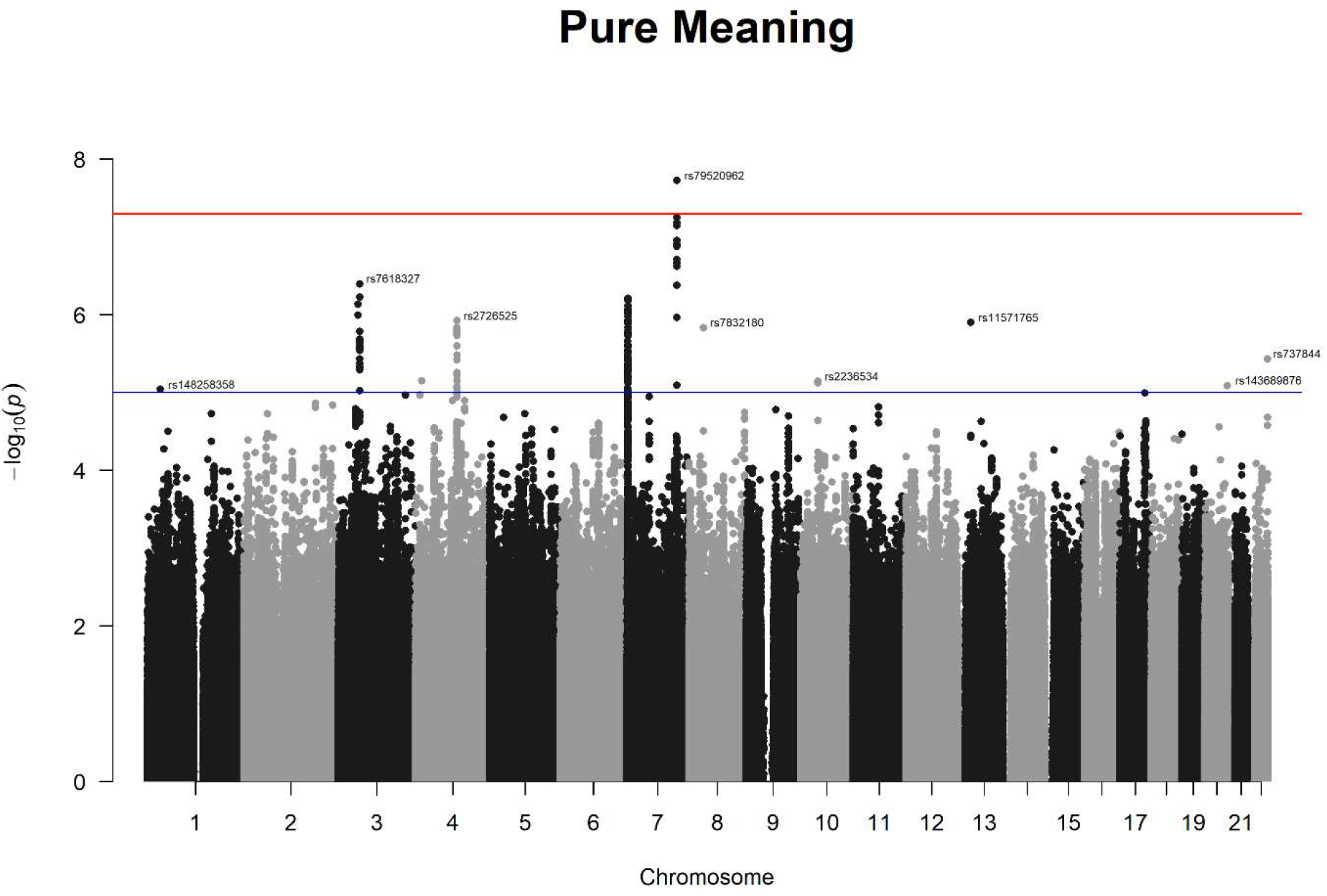
Manhattan plot for the GWAS results of pure meaning in life.

#### SNP heritability and genetic correlation

The SNP heritability of pure meaning was estimated to be 4.2% (*SE* = 0.005), a reduction of 2% compared to the SNP h^2^ of 6.2% (*SE* = 0.005) (5). The genetic correlation between pure meaning and meaning was 0.80 (*SE* = .04, *Z* = 18.36,*p* < .001), indicating a reduction in genetic (co)variance.

#### Functional annotation

The significant SNP rs79520962 (*β* = 0.304, *p* = 1.86×10^-8^) was also genome-wide significant in Baselmans and Bartels (2018) (*β* = 0.051, *p* = 2×10^-9^), with a similar effect size. The look-up showed no other associations for this SNP.

Applying FUMA, no gene replicated across the three different mapping methods. However, two genes, SND1 and LRRC4, were found through positional mapping, and SND1 was also found in the eQTL mapping. SND1 was also associated to meaning in life before the subtraction (5). The proteins encoded by SND1 are involved in cell growth. No genes were associated with pure meaning based on the gene-based tests and no significant enrichment of genes in certain tissues was found.

### Genetic correlations

The genetic correlation between pure happiness and pure meaning was estimated to be .65 (*SE* = .05, *p* = 1.25×10^-40^). Genetic correlations between happiness, meaning, pure happiness, pure meaning, and depressive symptoms can be found in supplementary Table S3.

The genetic correlations of pure happiness, pure meaning, happiness, meaning and depressive symptoms with all traits across 12 categories (N = 75) can be found in supplementary Figure S4 and Table S4. All correlations between the traits and respectively pure happiness and pure meaning had overlapping confidence intervals. Therefore, we refer to pure well-being instead of discussing the correlations separately for pure happiness and pure meaning.

In Figure 4 and 5, the genetic correlations with selected traits can be seen. We selected traits with a high correlation with well-being or depressive symptoms (Figure 4), and traits for which the genetic correlation with well-being versus pure well-being changed substantially or reversed (Figure 5).

**Figure 4.**
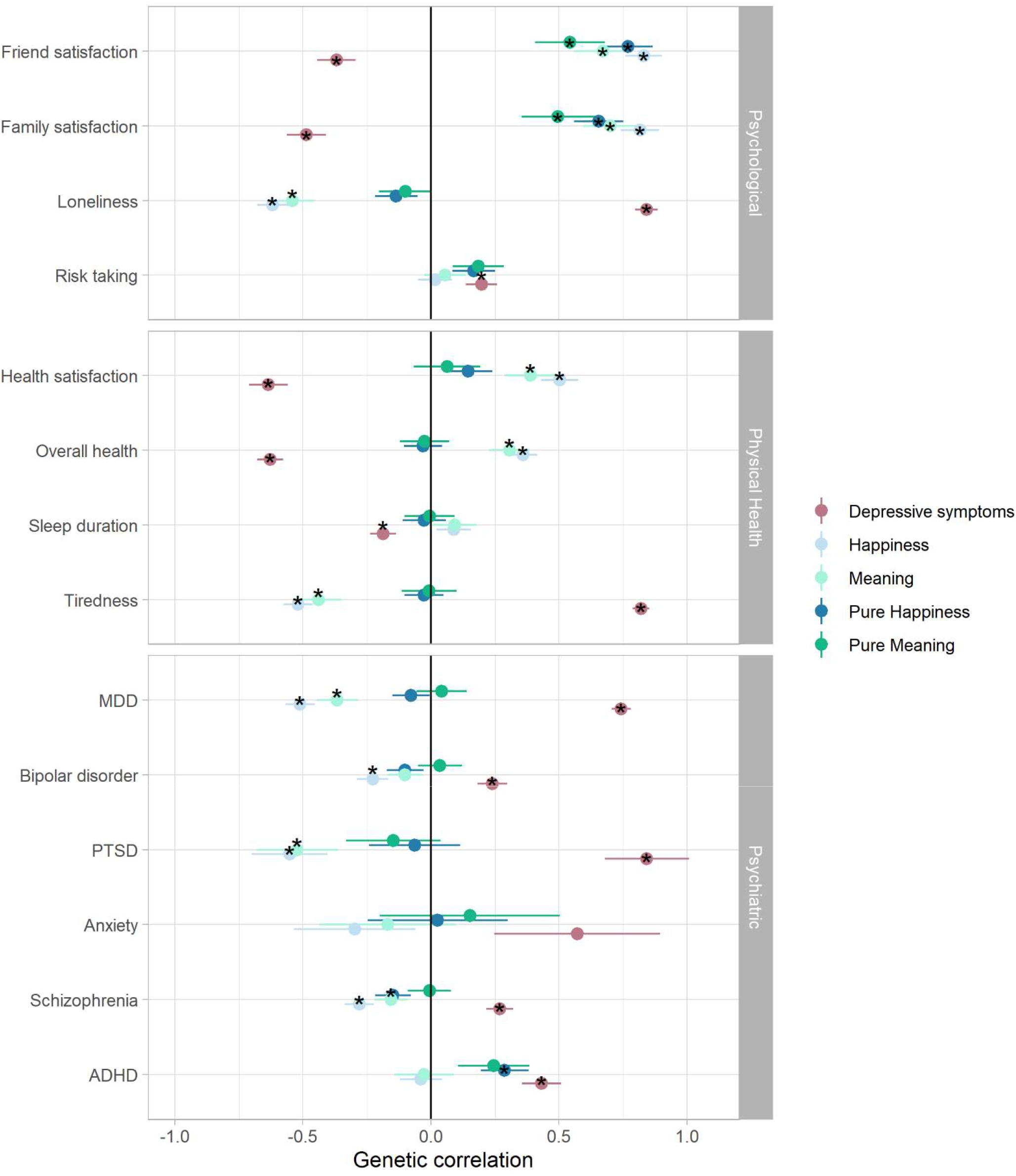
Genetic correlations of pure happiness, pure meaning, happiness, meaning and depressive symptoms and selected psychological traits, physical health traits, and psychiatric disorders. * indicates significant genetic correlations with a Bonferroni corrected threshold of *p* < 1.3×10^-5^.

**Figure 5.**
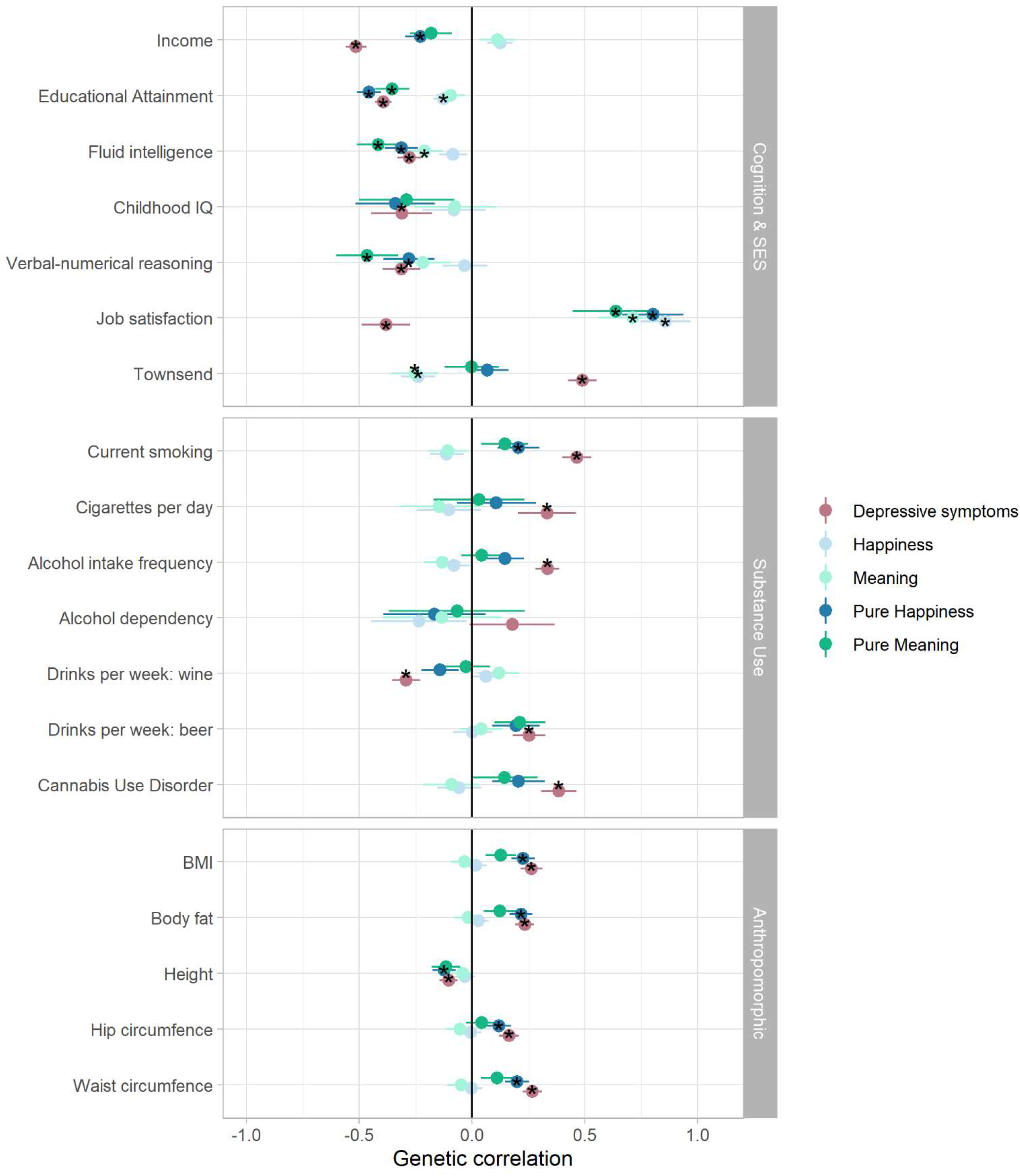
Genetic correlations of pure happiness, pure meaning, happiness, meaning and depressive symptoms and selected cognition and socio-economic status, substance use and anthropomorphic traits. * indicates significant genetic correlations with a Bonferroni corrected threshold of *p* < 1.3×10^-5^.

Three different patterns of genetic correlations were found. Different patterns of genetic correlations emerged, including (1) non-changing genetic correlations, (2) changed correlations from significant to zero, and (3) increased or reversed genetic correlations.

First, the subtraction of depressive symptoms did not influence the high genetic correlations of well-being with friend, family (Figure 4, psychological category), and job satisfaction (Figure 5).

Second, for psychological traits, psychiatric disorders and physical health traits related to depressive symptoms, pure well-being was not associated genetically. The subtraction of depressive symptoms GWAS removed the negative genetic correlations with well-being. This indicates that the original genetic correlations between well-being and depression-related traits are mostly due to the genetic overlap with depressive symptoms.

Third, genetic correlations for pure well-being versus well-being and several other traits increased or reversed (see Figure 5). For example, the genetic correlations between Attention deficit hyperactivity disorder (ADHD) and pure happiness and meaning (r_g_ = .29 and .25) became positive, compared to the non-significant correlations of well-being (r_g_ = −.04 and −.03). Similar results were found for risk taking. This indicates that a higher genetic predisposition for pure well-being is related to a higher genetic risk of ADHD and risk-taking, when corrected for the genetic predisposition for depressive symptoms.

Reversed effects were also found for SES traits (see Figure 5). Income was slightly positively associated with happiness and meaning in life (r_g_ = .12 and .11) before subtraction. Subtracting depressive symptoms from well-being, the genetic correlations became negative for pure happiness and meaning (r_g_ = −.23 and −.18). The genetic correlations between educational attainment and pure well-being became significantly negative (respectively r_g_ = −.46 and −.35 for pure happiness and pure meaning), compared to the smaller correlations (r_g_ = −.13 and −.09) before subtraction.

A consistent pattern of reversed genetic correlations between pure well-being and substance use traits, body fat, and BMI was also found, although not all correlations reached significance after correcting for multiple testing (see Figure 5). Before subtraction, these traits were genetically unrelated or slightly negatively associated with well-being (r_g_ between −.08 and −.13), whereas the association with pure well-being became positive (r_g_ between .05 and .23) after subtracting depressive symptoms.

## Discussion

Subtracting a depressive symptoms GWAS from happiness and meaning in life GWASs generated GWASs capturing genetic variants associated with happiness and meaning in life independent of depressive symptoms, i.e. ‘pure’ happiness and ‘pure’ meaning. For both latent traits, one independent SNP reached genome-wide significance (rs1078141 and rs79520962, respectively). Consistent with the larger genetic overlap of depression with happiness (r_g_ = −.53) compared to meaning in life (r_g_ = −.32) (5), we report a stronger reduction in SNP heritability of happiness (48%) compared to meaning in life (32%) after the subtraction of the depressive symptoms GWAS. The small reduction of the genetic correlation between happiness and meaning in life after the subtraction of depressive symptoms (r_g_=.78 to r_g_=.65) indicates that only part of the overlap between happiness and meaning in life is due to the overlap of the well-being measures with depressive symptoms. The largest part of genetic factors underlying happiness and meaning in life remains shared. Furthermore, the similar patterns of genetic correlations for pure happiness and pure meaning with a range of other traits are in line with a largely shared genetic etiology.

### Pure well-being correlates

The genetic correlations of well-being with other traits before and after the subtraction of depressive symptoms led to insights about the unique part of well-being. Different patterns of genetic correlations emerged, including (1) non-changing genetic correlations, (2) changed correlations from significant to zero, and (3) reversed genetic correlations. We discuss the meaning and implications of these different patterns of genetic correlations below.

First, genetic correlations between pure well-being and respectively family, friend, and job satisfaction did not change compared to the genetic correlations with well-being. The genetic predisposition to be satisfied with different life aspects is therefore related to the unique part of well-being and unrelated to the genetic predisposition for depressive symptoms. An exception is health satisfaction, being strongly related to depressive symptoms, the genetic correlation with pure well-being became non-significant, in line with the traits discussed next.

Second, as one could expect, pure well-being became genetically unrelated to traits correlating strongly with depressive symptoms, i.e., tiredness, overall health, and psychiatric disorders like post-traumatic stress disorder, bipolar disorder, and schizophrenia. This pattern indicates that part of the genetic variance of well-being can be seen on the continuum from depressive symptoms to well-being. The associations of well-being with these depression-related traits arise from the overlap with depressive symptoms and should be interpreted considering the current findings.

Third, for several other traits, including ADHD, SES, and substance use, the genetic correlations with well-being changed substantially after the subtraction of depressive symptoms. This indicates unique genetic overlap between pure well-being and these traits, independently from depressive symptoms. We shortly discuss possible explanations and mechanisms underlying the changed genetic correlations for these traits.

#### Attention-deficit/hyperactivity disorder (ADHD)

ADHD is a neurodevelopment disorder including symptoms of impaired attention, hyperactivity and impulsivity (23). ADHD is related to poor outcomes in academic achievement, and work performance (24–26). A positive genetic correlation between ADHD and depressive symptoms has been reported (e.g., see (27)). Our results indicate that a higher genetic predisposition for ADHD is also related to a higher genetic predisposition for pure wellbeing. An explanation for this finding could be the benefits and positive effects of ADHD symptoms in well-functioning individuals. Positive traits associated with ADHD include hyper focus, creativity, spontaneity, resilience, and high energy (28–31). These benefits are also related to well-being (32,33), suggesting the genetic correlation between pure well-being and ADHD captures the benefits of ADHD, when taking out the genetic predisposition for depressive symptoms.

#### Income, educational attainment (EA) and intelligence

Income, EA, and intelligence are strongly interrelated (13,34). We found similar effects of subtracting a depressive symptoms GWAS from well-being on the genetic correlations with these traits. The pure well-being genetic correlations indicated that people with a higher genetic predisposition for pure well-being also have a genetic predisposition for lower income, EA, and intelligence. Furthermore, the genetic correlations became similar in magnitude to the negative correlations of these traits with depression (35). Different mechanisms underlying the negative genetic correlation between depressive symptoms and income/EA/intelligence have been proposed. Low income/EA/intelligence can increase the risk of depression or vice versa, depressive symptoms have detrimental effects on the ability to actively and optimally participate in school and the labor force, leading to lower EA and incomes (36). The slightly positive genetic correlation between well-being and the traits before subtraction seems to be driven by the shared part with depressive symptoms, i.e., the opposite effects of depressive symptoms.

Possible explanations for the reversed genetic correlations for pure well-being can be nonlinear relations between well-being and income, EA, or intelligence. For example, for both income and intelligence, satiation and turning points on well-being have been found. The satiation point of income suggests that above a certain level of income that is sufficient to fulfil basic physical needs, higher income does not lead to higher levels of well-being (37). The turning point of income indicates that people with very high incomes report lower well-being levels compared to those with lower incomes (38). However, results tend to depend on analytic approaches (39). Similar, in highly intellectually “gifted” individuals (i.e., very high intellect, IQ ≥ 130), lower levels of well-being have been found compared to high-achieving individuals (i.e., only a high performance) and the general population (40,41). Intellectually gifted individuals can be at a greater risk for the development of a meaning in life crisis. To further explore the negative genetic associations between pure well-being and income/intelligence, we performed exploratory analyses on phenotypic associations between income/intelligence and happiness/meaning in life in a high and low income/intelligence group (see supplementary Material). The results were inconclusive. Therefore, and because the UK Biobank sample has a higher socioeconomic status compared to the general population, more research on the association between income/EA/intelligence, well-being, and depressive symptoms in a multivariate design is needed to test these relations.

#### Substance use and food-related traits

A consistent pattern of genetic correlations between pure well-being and substance use and food-related traits appeared as well, although not all correlations reached significance after correcting for multiple testing. Smoking, alcohol intake frequency, BMI, and body fat were genetically positively related to depressive symptoms and unrelated or slightly negatively associated to well-being. Pure well-being became positively genetically related to these traits after subtracting depressive symptoms. A possible explanation for these reversed genetic correlations for pure well-being could be the different underlying reasons why people smoke, drink, and eat. The genetic overlap between depressive symptoms and these traits can arise from self-medication, i.e., smoking, drinking and eating to cope and reduce the negative mood or other depressive symptoms (42–46). In contrast, smoking, drinking, and eating that is genetically related to pure well-being could arise from these behaviors in social settings. More research is needed to investigate the specific associations of well-being and substance use and eating variables.

### Limitations and implications

The results should be interpreted in light of some limitations. The UK Biobank sample is known to be biased, participants are on average older, healthier, include more females, and have a higher socioeconomic status compared to the general population (47). Therefore, subtraction in UK Biobank GWASs might have introduced extra bias in the pure well-being GWASs, possibly influencing the results and genetic correlations of pure well-being with other traits. Furthermore, UK Biobank focuses on samples from European ancestry. Well-being is differently conceptualized in different cultures (48), limiting generalization across samples with other ancestries. Replication of these results using GWASs from population-wide samples and more ancestry-diverse samples is needed.

If replicated, the findings can have important implications for mental health research and preventions or interventions. Note that we reported genetic correlations, indicating genetic sensitivity to both traits and we did not investigate direct phenotypic associations or causal effects. The patterns of genetic correlations of well-being versus pure well-being indicate that the genetic variance of well-being can be split into two parts having different associations with other traits. Part of the variance of well-being is overlapping with depressive symptoms, whereas the other part is unique to well-being. Based on our results, different associations could therefore be taken into account depending on the goal of the intervention. If the goal is to both decrease depressive symptoms and increase well-being, interventions should consider variables that are (causally) related to both depressive symptoms and well-being. If the goal is to increase well-being, instead of just reducing depressive symptoms, interventions should focus on variables that are (causally) related to pure well-being. Our results can be used as a starting point to find these variables and design future interventions.

## Supporting information

Supplemental Sensitivity Analyses + Figures

## Acknowledgements and disclosures

### Data availability

The data that support the findings are available via the UK Biobank. The preregistered analytical plan is shared on OSF (https://osf.io/pnc2z).

### Funding

This work is supported by an ERC consolidation grant (WELL-BEING 771057 PI Bartels) and by the grant 531003014 from The Netherlands Organisation for Health Research and Development (ZonMW).

### Ethics approval and consent to participate

All procedures performed in studies involving human participants were in accordance with the ethical standards of the institutional and/or national research committee and with the 1964 Helsinki declaration. Informed consent was obtained from all individual participants included in the study.

## Acknowledgements

The authors would like to thank all participants, who provided data for this study.

## Conflict of interest

The author(s) declared that there were no competing interest with respect to the authorship or the publication of this article.

## Notes

### Competing Interest Statement

The authors have declared no competing interest.

