## Supplemental Sensitivity Analyses + Figures for "Distinguishing Happiness and Meaning in Life from Depressive Symptoms: a GWAS-by-subtraction study in the UK Biobank"

**Supplementary materials**

**Sensitivity** **analysis**

The GWAS-by-subtraction model assumes that the genetic effects on depressive symptoms have an effect on happiness or meaning in life. There could be a possible bidirectional effect between depressive symptoms and well-being. The possible bidirectional causal effect can be considered a violation of the assumption. To investigate the impact of a bidirectional effect between depressive symptoms and well-being, we allowed for this effect in the model (see supplementary Figure S5 for the adjusted model). We cannot estimate the effect freely, because of identification issues and therefore included a small effect of 0.2. Including this small effect of well-being on depressive symptoms, we reanalyzed and investigated the change in the Z-statistics of the genome-wide and suggestive SNPs for happiness and meaning in life. There was a minimal change in the Z-statistics, both for pure happiness and pure meaning (see supplementary Figure S6 for the comparison in Z-statistics).

**Phenotypic correlations**

To further explore the negative genetic associations between pure well-being and income, we did an exploratory phenotypic analysis and compared the phenotypic correlations between income and happiness/meaning in life in a high and low income group (median 50% split). In contrast to these suggestions of the highest incomes to be related to lower well-being, we found a small positive correlation between income and respectively happiness and meaning in life for both the high (r_ph_ = .05 and .04) and low income group (r_ph_= .08 and .06). A limitation of this analysis is that the UK Biobank sample has a relatively higher income compared to the general population. Furthermore, the relation between lower well-being and income is suggested to only occur in the highest incomes, probably not captured by the median split. Finally, we did not take out the effects of depressive symptoms on income in the phenotypic correlations. Therefore, more research on the association between income, well-being, and depressive symptoms in a multivariate design and in appropriate samples is needed to test the associations.

Similarly, as exploratory analysis, we compared the phenotypic correlations between intelligence and happiness/meaning in life in a high and low intelligence group (median 50% split). Although the correlations are very small, we found small phenotypic correlations in line with these ideas. For happiness, we found a positive correlation between income and happiness in the low intelligence group (r_ph_= .03) and a small negative correlation in the high intelligence group (r_ph_= -.01). For meaning in life, the negative correlation with intelligence was stronger in the high intelligence group (r_ph_= -.04) compared to the low intelligence group (r_ph_=-.01). However, as the UK Biobank sample has a higher socioeconomic status compared to the general population, we did not take depressive symptoms into account, and the phenotypic correlations are very small, more research on the (phenotypic) associations between intelligence, well-being and depressive symptoms is needed to test these hypotheses.

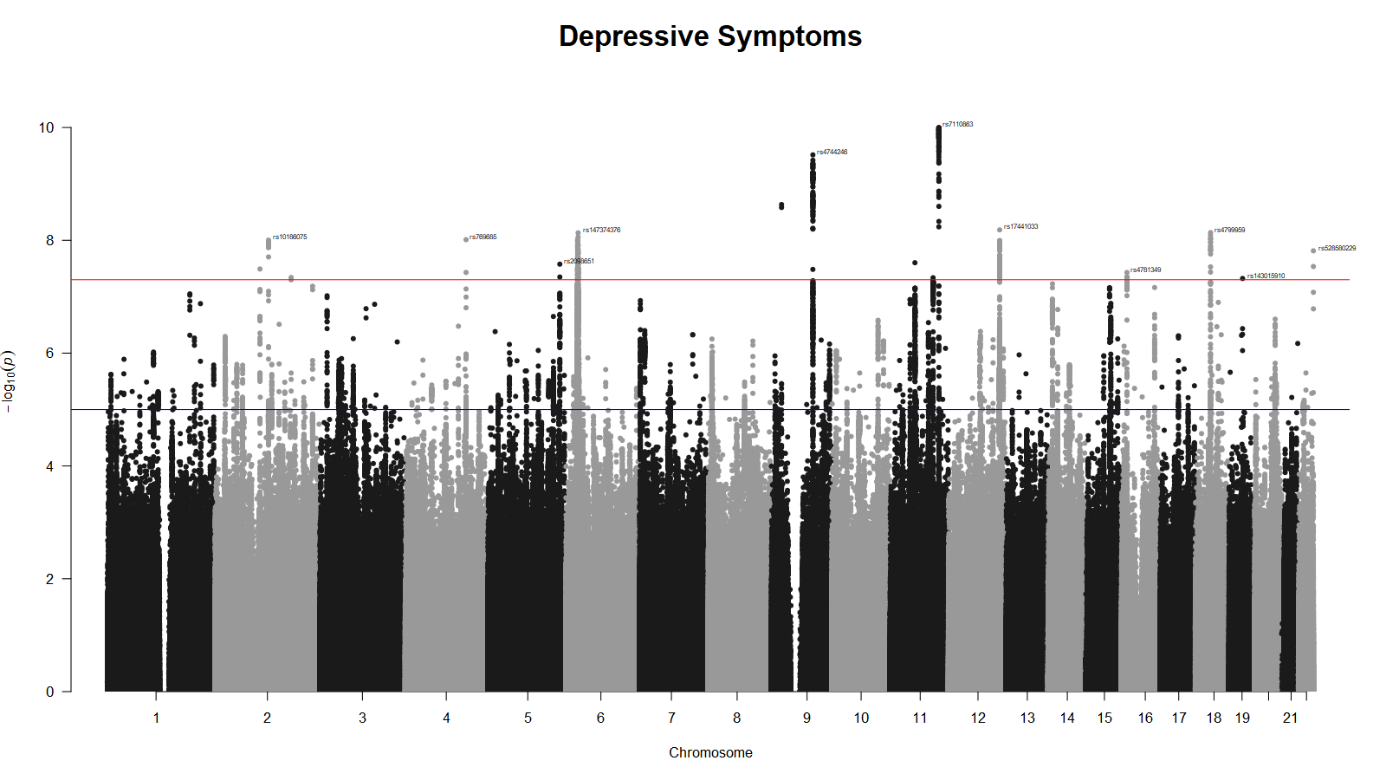

**Supplementary Figure S1.** Manhattan plot for the GWAS results of depressive symptoms.

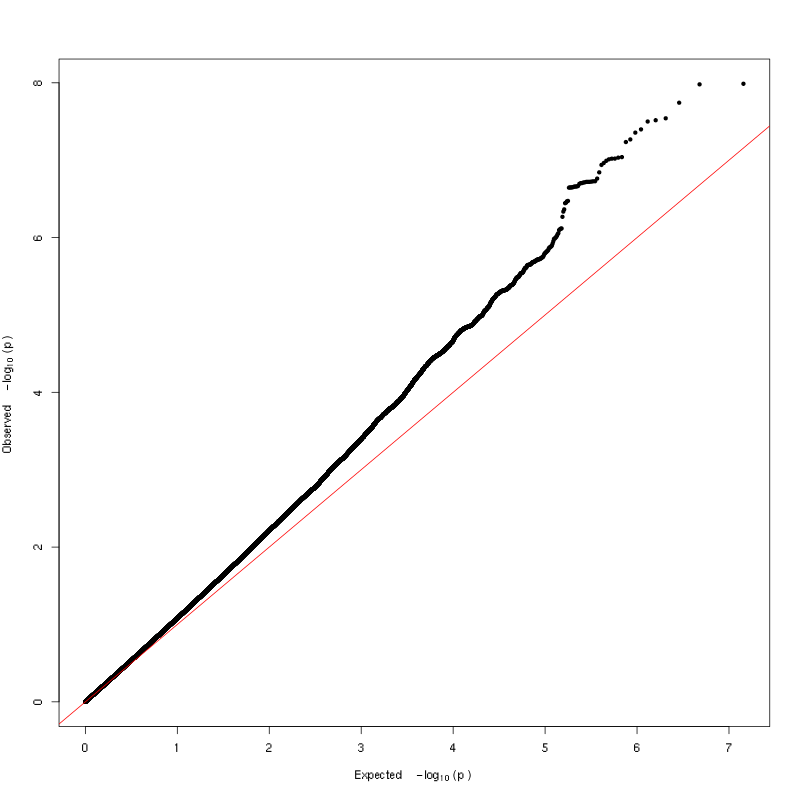

**Supplementary Figure S2.** QQ plot for pure happiness GWAS.

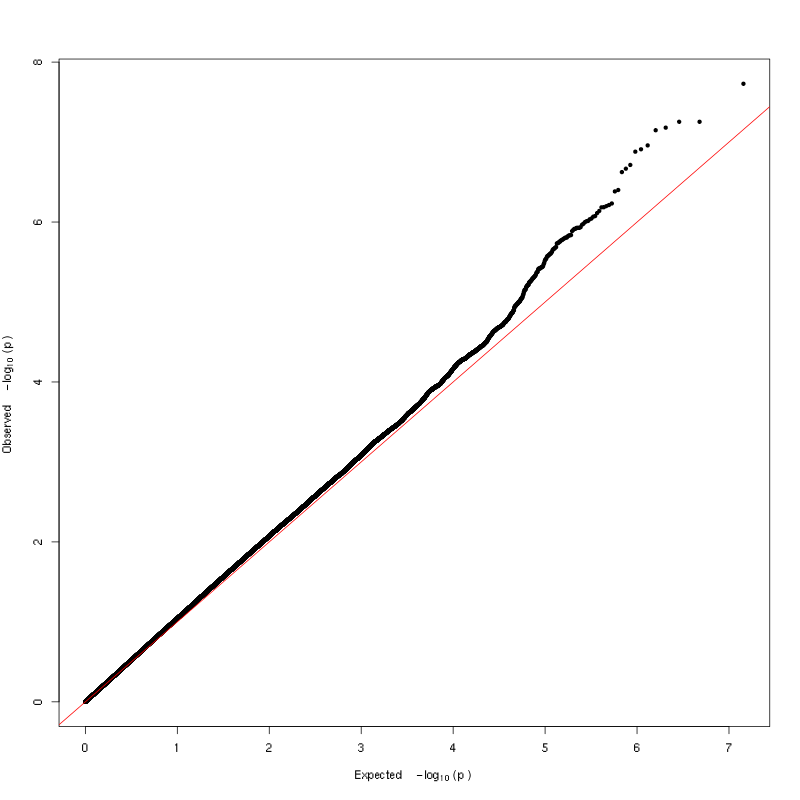

**Supplementary Figure S3.** QQ plot for pure meaning GWAS.

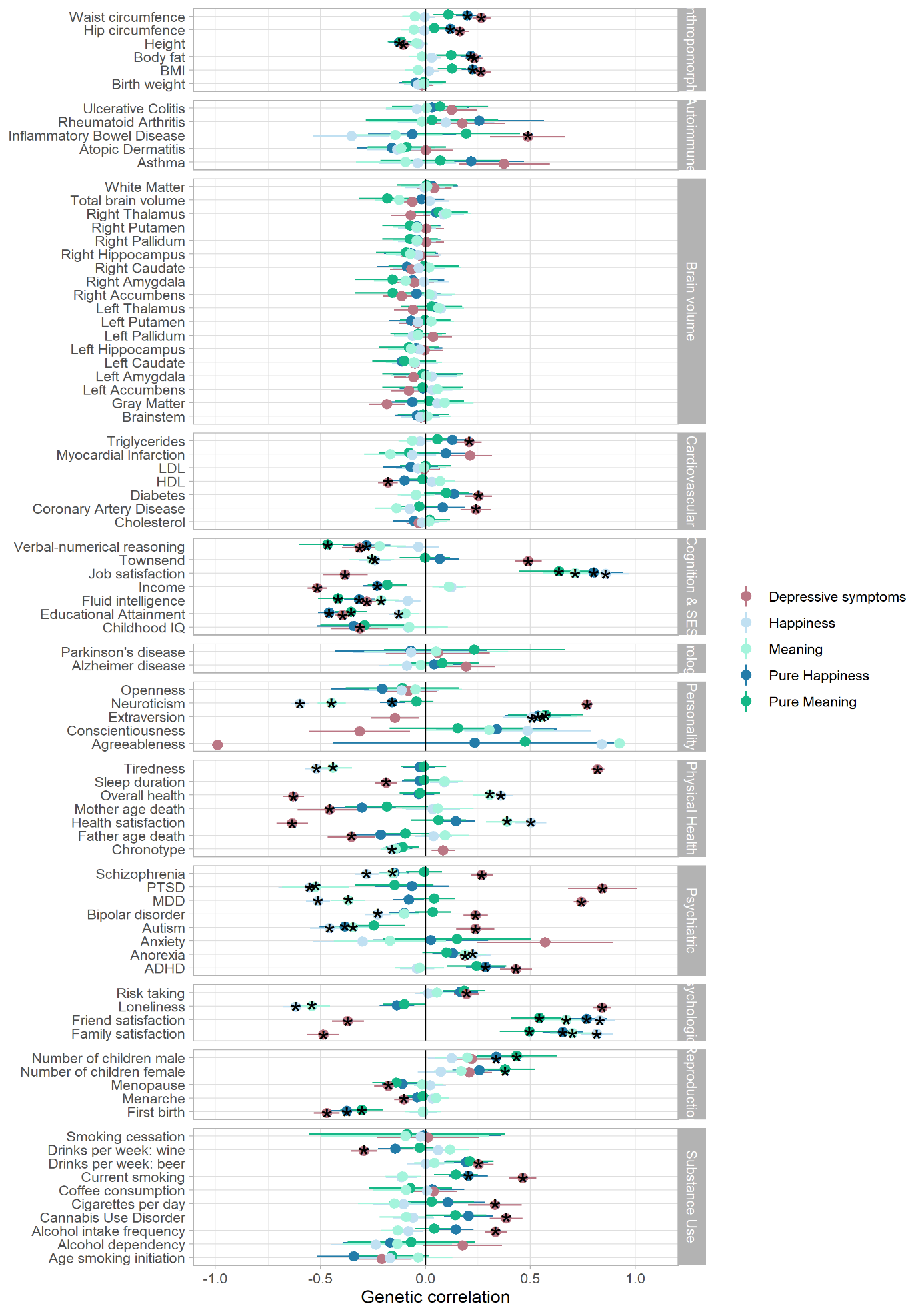
 **Supplementary Figure S4.** Genetic correlations between happiness, meaning, depressive symptoms, pure happiness and pure meaning and a range of different traits across 12 categories (n=75). Stars indicate significant correlations, Bonferroni corrected (*p*=.00013).

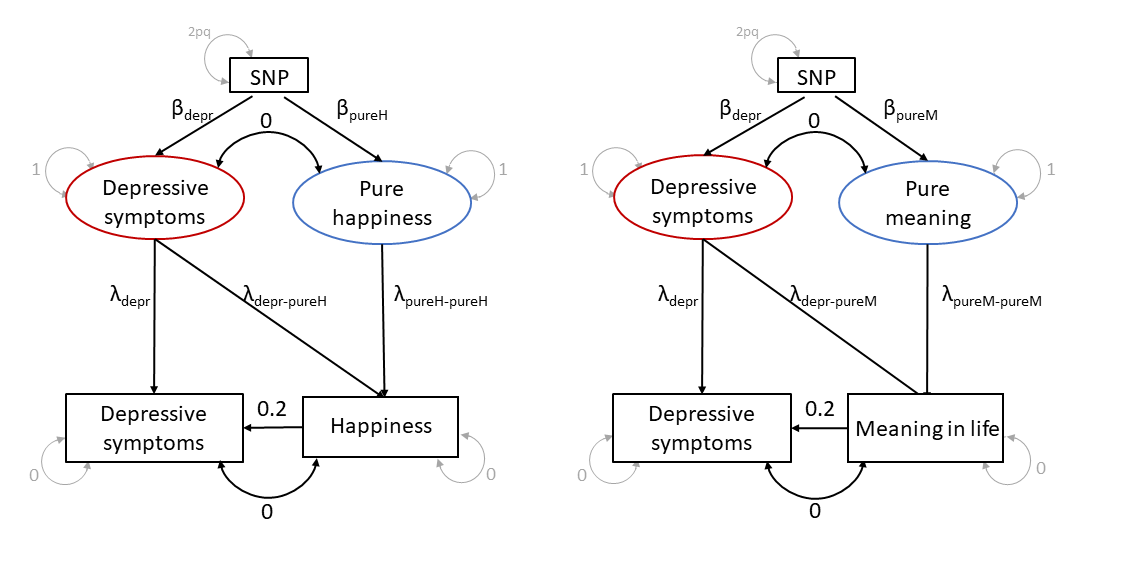

**Supplementary Figure S5.** GWAS-by-subtraction model with a small reversed causal effect of 0.2 of happiness (left panel) and meaning in life (right panel) on depressive symptoms.

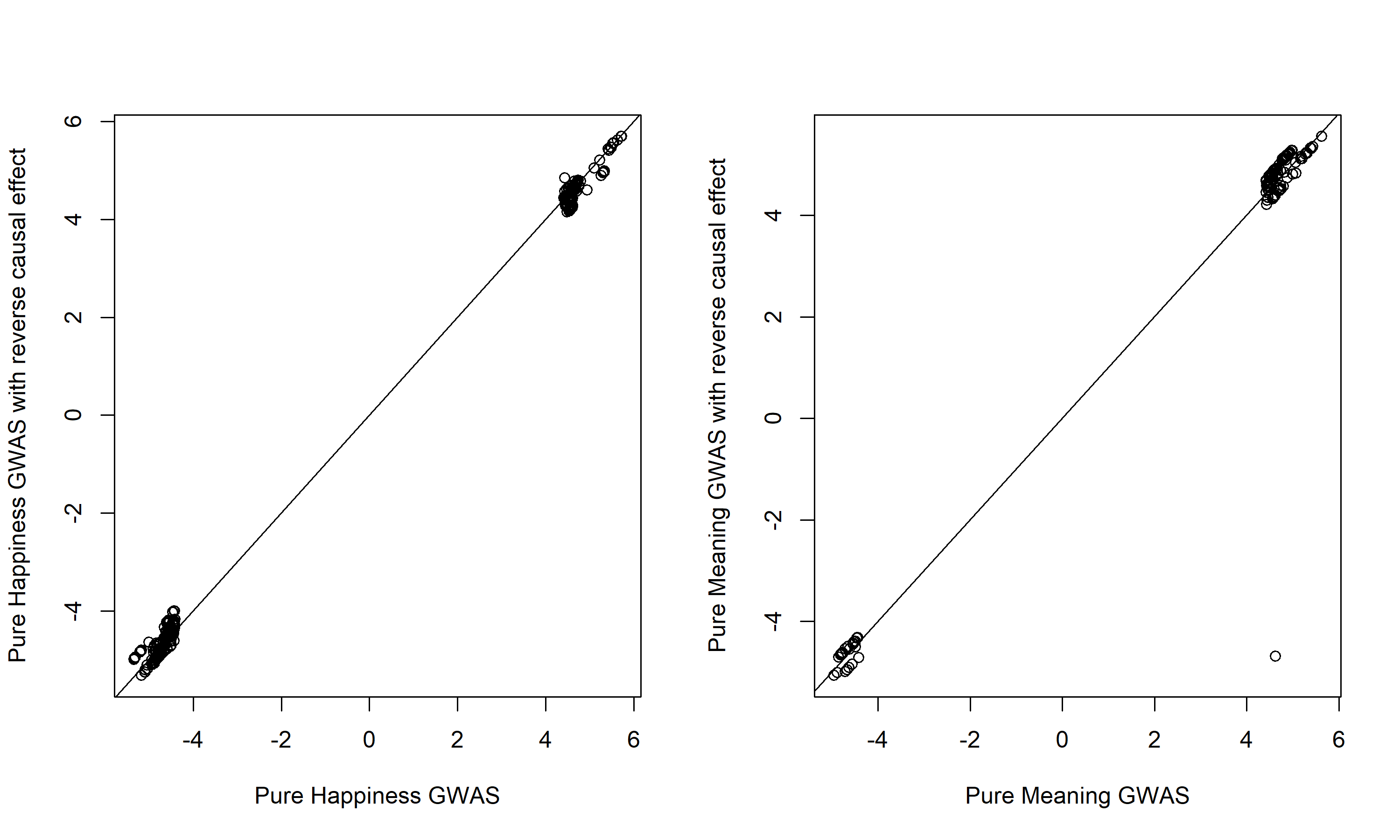

**Supplementary Figure S6.** Comparison of the Z-statistics of the hits and suggestive SNPs estimated in the standard GWAS-by-subtraction model and in a model with a small reversed causal effect (0.2) from happiness (left panel: 345 SNPs) and meaning in life (right panel: 138 SNPs) to depressive symptoms.
